# A scalable organoid model of urothelial aging for metabolic interrogation, infection modeling, and reversal of age-associated changes

**DOI:** 10.1101/2025.06.27.662009

**Authors:** Adwaita R. Parab, Arnold M. Salazar, Steven J. Bark, Margarita Divenko, Vasanta Putluri, D’Feau Lieu, Aadya S. Singh, Nagireddy Putluri, Indira U. Mysorekar

## Abstract

Aging leads to a progressive decline in overall bladder function resulting in lower urinary tract symptoms and increased susceptibility to infections. However, tissue-specific mechanisms of aging, specifically the contributions of the aged urothelium remain elusive. Here, we introduce mouse bladder epithelium-derived organoids (mBEDOs) as a scalable platform to model urothelial aging. mBEDOs from aged mice recapitulate key features of age-associated cellular reprogramming, including oxidative stress, senescence, and DNA damage. We demonstrate the utility of mBEDOs for modeling Uropathogenic *Escherichia coli* (UPEC) infection, generating assembloids between mBEDOs and macrophages to model epithelial-immune interactions, and genetic perturbation. Using the mBEDO platform, we also identify urothelium-specific changes in purine, amino acid, and glycerophospholipid metabolism which may contribute to age-associated cellular perturbations. Lastly, supplementation with depleted metabolites, nicotinamide (NAM) and D-mannose, reduce DNA damage and oxidative stress and restore mitochondrial integrity in aged mBEDOs. These findings establish mBEDOs as an effective platform for investigating molecular and cellular underpinnings of urothelial aging and exploring metabolism-based interventions for age-associated bladder dysfunction.

## 1 Introduction

Aging is a principal driver of disease, characterized by diminished cellular homeostasis, impaired tissue repair, and increased susceptibility to infection(López-Otín et al., 2013). Age-associated changes are especially detrimental in epithelial barrier tissues, which serve as the first line of defense against environmental insults. In the lower urinary tract, age-associated effects include reduced bladder capacity, overactive bladder, decreased urethral pressure, and higher levels of residual urine in the bladder (Nishii, 2021; J. W. Kim et al., 2020). These functional issues manifest in the form of lower urinary tract symptoms (LUTS) such as urinary incontinence, lack of voiding control, bladder pain syndrome, and recurrent urinary tract infections (rUTIs)(Hu et al., 2004; Coyne et al., 2009; Heidler et al., 2007; Tsai et al., 2018).

The bladder epithelium, or urothelium, plays a vital role in urinary tract health by forming a barrier against toxins and pathogens and responding to mechanical stressors such as stretch and osmotic pressure (Jafari & Rohn, 2022). Our previous work has shown that aged female mice (15–24 months) exhibit several hallmarks of urothelial aging, including increased senescence, spontaneous pyroptosis, oxidative stress, mitochondrial dysfunction, and impaired autophagy with lysosomal accumulation, which overall heighten UTI susceptibility (Joshi et al., 2024). In parallel, aging also affects bladder mucosal immunity. Bladders, like other mucosal tissues, also contain a diverse population of resident immune cells, with macrophages, the most abundant subset, forming a dense network beneath the urothelium, regulating tissue homeostasis and epithelial barrier maintenance (Lacerda Mariano & Ingersoll, 2018; Lacerda Mariano & Ingersoll, 2020). However, with age, B and T cells in addition to macrophages accumulate and organize into bladder tertiary lymphoid tissue (bTLT), structures that are rarely seen in healthy young but become prominent in aged female mouse bladders (Ligon et al., 2020). In the bladder, examining urothelium-specific aging phenotypes *in vivo* is complicated by interactions with various components such as immune cells, vasculature, neurons, and smooth muscle. This gap highlights the need for model systems that can be used to study epithelial-intrinsic changes in isolation while also enabling study of epithelial-immune interactions.

Human and rodent models have been instrumental in advancing our understanding of aging and bladder biology. However, studying the functional consequences of aging in a tissue-specific manner, particularly through genetic perturbation, remains a significant challenge. Performing loss-of-function studies in aged mice is time-intensive, costly, and often not feasible at scale, as it requires aging colonies over many months and coordinating treatment windows late in life (McLean et al., 1992). Thus, complementary models that retain the strengths of mouse biology while enabling faster, more tractable interrogation of aging mechanisms in the urothelium are urgently needed.

The earliest studies describing *ex vivo* urothelial cell cultures date back to the 2000s when *ex vivo* urothelial constructs expressing uroplakins and cytokeratin 20, and retaining barrier characteristics were developed (Sugasi et al., 2000; Shin et al., 2011; Varley & Southgate, 2011; Vasyutin et al., 2019). Human urothelial cell monolayer cultures were developed where a stratified urothelium was generated from human bladder washings (Nagele et al., 2008). Recently, a urine-tolerant urothelial organoid model derived from commercially available human progenitors has been developed and shown to successfully model infection (Horsley et al., 2018; Smith et al., 2006). Another study showed interaction between the immune and epithelial compartments during infection, using a 3D mouse urothelial organoid model to show early bacterial invasion by UPEC and neutrophil response in an *ex vivo* system(Sharma et al., 2021). Additionally, studies have shown urothelial organoids derived from both mouse and human tissue to study bladder cancer (Mullenders et al., 2019; E. Kim et al., 2020; Mullenders et al., 2019) and use of bladder assembloids with stromal components and muscle layer to recapitulate the *in vivo* tumor microenvironment (E. Kim et al., 2020). However, organoid models to investigate aging-associated molecular, cellular, and metabolic alterations have not been developed.

In this report, we describe the development of mouse bladder epithelium-derived organoids (mBEDOs) as a scalable, tissue-specific model of urothelial aging. We demonstrate that mBEDOs from aged mice recapitulate key features of *in vivo* urothelial aging, including elevated oxidative stress, DNA damage, and cellular senescence. mBEDOs can be cultured in both 2D and 3D formats and extended into assembloids, incorporating immune cells, enabling modeling of uropathogenic *Escherichia coli* (UPEC) infection and epithelial–immune interactions. Importantly, mBEDOs can also be generated from genetically modified mice, offering a unique platform to examine the role of specific genes (Irg1) in the context of epithelial aging. Metabolomic characterization of young and aged mBEDOs and those generated from *Irg1^-/-^* mice using untargeted LC-MS/MS metabolomics identified distinct metabolic signatures of age. Finally, supplementing aged mBEDOs with D-mannose and nicotinamide (NAM), two metabolites depleted in aged samples, significantly reduced oxidative stress and DNA damage while restoring mitochondrial integrity. Together, these findings position mBEDOs and assembloids as powerful tools for dissecting epithelial aging, modeling infection susceptibility, and evaluating metabolism-based interventions in a high-throughput, tissue-specific context.

## 2 Methods

### 2.1 Organoid generation from mouse bladders

C57B6/J female mice (8-12 weeks: young; and 15-18 months: aged) were sacrificed and their bladders were extracted. Whole bladders were rinsed in cold PBS and minced into smaller pieces using a scalpel. Minced bladder tissue from each mouse was transferred to a 15 mL tube containing 1 mL of 5 mg/mL Collagenase II (Gibco Cat. No. 17101015), 0.5 µL 10 mM Y-27632 (Rock inhibitor) (Stem Cell Technologies Cat. No. 72302), 20 µL 10 mg/mL Elastase (Sigma-Aldrich Cat. No. E7885) and were placed in the Roto-Therm-Mini Mix Plus at 37°C and 10 RPM for four hours. The tubes were then centrifuged at 4°C and 1500 RCF for six minutes. The supernatants were aspirated, and the pellet resuspended in 1mL warmed TrypLE. The tubes were then placed in the Roto-therm-Mini Mix Plus at 37°C and 10 RPM for 20 minutes. Tubes were centrifuged again at 4°C and 1500 RCF for six minutes. The supernatants discarded and the pellet was resuspended in 1 mL warmed Advanced DMEM (Invitrogen (Cat. No. 12634-034) with GlutaMAX (Gibco Cat No. 350561). The cells were counted, and 20,000 cells were plated in 50 μL matrigel tabs in a pre-warmed 6-well suspension plate. The plate was then placed at 37°C for 15 minutes and added with 2 mL warmed proliferation medium (Advanced DMEM (Invitrogen Cat. No. 12634-034) supplemented with B27 (Invitrogen Cat. No. 12634-034), N-Acetyl-L-Cysteine (Sigma-Aldrich Cat. No. A9165-5G), NAM (Sigma-Aldrich (Cat. No. N0636-100G), mouse EGF (Life Technologies Cat. No. PMG8041), A-83-01 (Tocris Cat. No. 2939), 2% Noggin and 10% R-Spondin (both obtained from the Digestive Diseases Core at Baylor College of Medicine), 1.25% GlutaMAX (Invitrogen Cat. No. 12634-034), and 1% Pen Strep (Invitrogen (Cat. No. 15140-122). Medium was refreshed every two days. After seven days, medium was replaced with intermediate medium consisting of modified proliferation medium to include 12% WNT-3A (obtained from the Digestive Diseases Core at Baylor College of Medicine) and 10 pg/mL mouse IL-6 (Cell Signaling Technologies Cat No. 5216). On day 14, organoids were placed in differentiation medium (Advanced DMEM supplemented with 20% FBS, B27, A83-01, 25ng/mL mouse FGF-7 (PeproTech Cat. No. 450-60) and 100 ng/mL mouse FGF-10 (PeproTech Cat. No. 450-61) for seven days. Organoids were able to be maintained in media containing 10%FBS, Pen Strep in Advanced DMEM for about 10 days post-maturation.

### 2.2 Organoid passaging

Organoids were subcultured at the end of proliferation stage to enable multiple technical replications. Organoids were successfully subcultured up to 10 passages. To subculture organoids, organoids in matrigel tabs were gently disrupted using a p1000 pipette tip in the medium. Organoid suspension from each well were transferred to a 15 mL tube and centrifuged at 4°C and 1500 RCF for six minutes. Supernatant was discarded and cells were resuspended in 1mL ice-cold PBS and incubated on ice for 10 minutes. Tubes were then centrifuged at 4°C and 1500 RCF for six minutes. This was repeated two more times to remove most of the Matrigel from the organoid pellet. The pellet was then resuspended in 1 mL warmed Advanced DMEM. Cells were counted and 20,000 cells were placed in 50 μL matrigel tabs in warmed 6-well plates. Freezer stocks with 1×10^6^ cells were made by resuspending organoids in Recovery Cell Culture Freezing Medium (Gibco Cat No. 12648010) and stored in liquid nitrogen for future passages.

### 2.3 Organoid preparation for immunohistochemistry and histology

6-well plates with organoids were incubated on ice for 10 minutes. The 1000 µL pipette tip was cut diagonally and used to gently break the Matrigel in the medium. The organoid suspensions were collected and transferred to a 15 mL tube. The tube was then incubated on ice for 10 minutes following which it was centrifuged at 1500 RCF for five minutes at 4°C. The supernatant was discarded, and the pellet was gently resuspended in ice-cold PBS. The tube was centrifuged, and the supernatant was discarded. The PBS washes were repeated twice, and organoids were transferred to a 1.5 mL tube. OCT cassettes were prepared by adding a thin layer of pelleted organoids in OCT, followed by freezing at −20°C and stored at-80°C

### 2.4 Immunofluorescence staining

Formalin-fixed paraffin-embedded organoid sections from young and aged mice were deparaffinized by soaking in 3 separate solutions of 100% Histoclear at 5 minutes each. Sections were rehydrated in decreasing concentrations of ethanol at 100%, 90%, 70% and 50% for 5 minutes at each concentration. Slides were then soaked in 1X PBS for 5 minutes to remove excess alcohol. 1% bovine serum albumin (BSA) in 1x PBS was used to block the slides for 1 hour at room temperature. Antibody staining was performed using anti-UPKIIIA (Fitzgerald Cat No. 10R-U103a), anti-CK5 (Abcam Cat No. ab53121), and anti-γH2AX (Cell SignalingTechnology Cat No. 9718S) in 1% BSA and 0.1% Tween-20 overnight at 4°C. Thereafter, slides were washed with 1X PBS 3 times for 5 minutes each, and incubated with the secondary antibody (Invitrogen Alexa Fluor Goat anti-Mouse 488 Cat No. A11029, Goat anti-Rabbit 594, Cat No. A11037 at 1:1000) for 1 hour at room temperature. After this, slides were washed again with 1X PBS, 3 times for 5 minutes each. 10 µL Prolong Gold Antifade reagent with DAPI (P36935, Thermo Fisher Scientific, USA) was added to each slide on the side with the section. The slides were covered with a cover slip and the edges were sealed with clear nail polish. Images were captured using a Nikon Ti2 ECLIPSE confocal microscope at 40X magnification, and 1.4 aperture. Fluorescence intensity was quantified from 3 mBEDO batches with 5 ROIs and further analyzed using the Nikon NIS Elements software.

### 2.5 H&E staining

Frozen OCT sections were fixed with 4% PFA for 10 minutes followed by rinsing with PBS three times for 3 minutes. Samples were rehydrated in decreasing concentrations of ethanol (100%, 95%, 70%, and 50% EtOH) for 5 minutes at each concentration. Excess alcohol was removed by rinsing with distilled water. Slides were dipped in Hematoxylin for 30 seconds. Excess hematoxylin was washed off with water followed by dipping in acid alcohol. Slides were then soaked in sodium bicarbonate for 3 minutes, followed by soaking in 90% ethanol. Slides were then dipped in Eosin 5 times. Following this, they were placed in increasing concentrations (50%, 70%, 95%, 100%) of ethanol for 5 minutes at each concentration. Finally, slides were soaked in Histoclear for 2 minutes, added with Permount mounting medium, and then cover-slipped. Images were captured using a Paranomic Midi microscope (3DHISTECH Ltd, Hungary).

### 2.6 Real-time quantitative PCR (RT-qPCR)

RNA was extracted from frozen pellets from young, aged, *Irg1^-/-^*mBEDOs by homogenizing with TRIzol™ Reagent (Thermo Fisher Scientific, 15596026, USA), and precipitated using chloroform. Samples separated into three layers where the top queous layer was RNA, middle cloudy layer was DNA, and bottom layer was protein. The top aqueous layer was carefully removed and RNA was precipitated with isopropanol. RNA was then washed with 75% ethanol and then the tubes were allowed to dry to obtain a RNA pellet. The RNA pellet was then solubilized in molecular biology grade water followed by DNase I treatment (Thermo Fisher Scientific, 18068-015, USA) according to the manufacturer’s instructions. 100 nanograms of Total RNA was used to synthesize cDNA with SuperScript™ II Reverse Transcriptase (Thermo Fisher Scientific, 18064-014, USA). All cDNA samples were diluted 1:8 with RNase-free water. Primers (**Supplemental Table 1**) and RT-qPCR reactions were prepared using SsoAdvanced Universal SYBR™ Green Supermix (Bio-Rad, 1725274, USA) in 10 µL reactions (5 µL SYBR, 1 µL each of Forward and Reverse Primers, 2 µL diluted cDNA, 1 µL RNase-free water). Reactions were performed in triplicate, with GAPDH as the reference gene. RT-qPCR was conducted on a QuantStudio™ 3 Real-Time PCR System (Applied Biosystems, USA) under the following conditions: 98°C for 3 minutes (initial activation), 98°C for 30 seconds (denaturation), 58°C for 30 seconds (annealing/extension), for 40 cycles. Raw Ct values were used to calculate fold changes relative to the young mBEDO samples. Final graphs were made and statistical analysis done using GraphPad Prism.

### 2.7 Derivation of 2D monolayers from 3D organoids

Matrigel tabs were disrupted to collect whole 3D organoids into 15 mL tubes. Tubes were centrifuged at 1500 RCF, 4°C for 5 minutes and then supernatants removed. 1 mL TrypLE was added to each tube and tubes were rotated at 37°C for 20 minutes. Tubes were centrifuged and supernatants removed. The organoid pellet was then resuspended in 20 µL of 1:20 Matrigel in proliferation media. Each 200 µL single-cell suspension was then plated in a well of a 24-well confocal plate. Media was changed every day for 4 days, after which a monolayer of urothelial cells was formed. This could be further sub-cultured or used for further experimentation.

### 2.8 UPEC infection of 2D monolayers

GFP-tagged UPEC was grown in static culture 75 mL LB in 250 ml flask at 37°C for 17-18h. Bacteria were centrifuged at 2000 rpm for 10 minutes. Bacteria was OD-adjusted to 0.5 at an absorbance of 600 to have 4×10^8^ *E. Coli* cells /ml in PBS. Monolayers were infected at MOI 1:100 with 50 µL OD adjusted *E. coli* in 950 µL antibiotic free Advanced DMEM. Monolayers were incubated with *E. coli* for 1 and 3 hours post-infection after which cells were washed with PBS and the antibiotic-free media was replaced with gentamicin (10mg/mL stock concentration) in 1:500 dilution in media.

### 2.9 Monolayer CFU analysis

Infected monolayers were harvested by scraping the wells of the plate using a 1000 µL pipette tip and transferring to a 1.5 mL tube. The cell lysate was centrifuged for 30 seconds and the supernatant collected. Serial dilutions of the supernatant were made in sterile PBS and 20 µL of each dilution was plated on an LB agar plate. 5 colony forming unit (CFUs) readings were collected per sample and averaged to obtain CFU readings which were log_10_ transformed and plotted using GraphPad Prism.

### 2.10 Western Blotting (WB)

mBEDOs were collected and stored in RIPA lysis buffer at −80°C. Lysis was done using sonication (each sample was sonicated 3 times for 3 seconds at 5 second intervals) and total protein was quantitated using a BCA assay (Pierce BCA Protein Assay Kit, 23225, Thermo Fisher Scientific, USA). 10µg of protein was loaded onto precast gel (4561095, 4–20% Mini-PROTEAN TGX Precast Protein Gels, Bio-Rad, USA) and resolved at 200 volts for 30-35 minutes. Immobilization of protein bands was done by transferring to PVDF membrane (IPFL00010, Immobilon Transfer Membrane, Millipore, Ireland) for one hour at 110 volts in ice. Blocking buffer (927-60001, Intercept (TBS) Blocking Buffer, LI-COR, USA) was used to treat the membrane for 1 hour at room temperature with gentle agitation. Membranes were incubated with primary antibody solution in blocking buffer plus 0.1% Tween-20, overnight at 4°C with agitation. Beta-actin was used as a loading control. Membranes were washed in 1X TBS with 0.1% Tween-20, 5 times, for 5 minutes each at room temperature with agitation followed by treatment with appropriate secondary antibody solution in blocking buffer plus 0.1% Tween-20 for 1 hour at room temperature with agitation. Membranes were washed in 1X TBS with 0.1% Tween-20, 5 times, for 5 minutes each at room temperature with agitation followed by washing in 1x TBS 2 times, for 5 minutes each. Imaging was performed using the ChemiDocTM MP imaging system (Bio-Rad, USA). Individual bands were quantified using Bio-Rad Image Lab software (6.0.1).

### 2.11 Lysosome Staining

Fresh-frozen 10 µm sections of young and aged mBEDOs were stained for lysosomes using Lysotracker Red dye. The sections were incubated in 75 nM Lysotracker dye solution for 30 minutes at 37°C, rinsed with 1X PBS and counterstained with DAPI. Sections were then mounted with Prolong Gold Antifade mountant (P10144, Thermo Fisher Scientific, USA), covered with cover slip, and the edges sealed with clear nail polish. Images were captured using a Nikon Ti2 ECLIPSE confocal microscope at 40X magnification, and 1.4 aperture, and further analyzed using the Nikon NIS Elements software. Lysosomes were quantified using 3 different mBEDO samples, and 5 ROIs were averaged to obtain mean fluorescence intensity in ImageJ software.

### 2.12 Mitochondria Staining

Fresh-frozen 10 µm sections of young and aged mBEDOs were stained for mitochondria using MitoTracker™ Orange CMTMRos and MitoTracker™ Green FM. The sections were incubated in 100 nM Mitotracker dye solution for 30 minutes at 37°C, rinsed with 1X PBS and counterstained with DAPI. After staining, the sections were mounted with Prolong Gold Antifade mountant (P10144, Thermo Fisher Scientific, USA), covered with a cover slip, and the edges sealed with clear nail polish. Images were captured using a Nikon Ti2 ECLIPSE confocal microscope at 40X magnification, 1.4 aperture, and further analyzed using the Nikon NIS Elements software. Mitochondria were quantified using 3 different mBEDO samples, and 5 ROIs were averaged to obtain mean fluorescence intensity in ImageJ software.

### 2.13 Dihydroethidium staining

Fresh-frozen 10 µm sections of young and aged mBEDOs were stained for reactive oxygen species using Dihydroethidium (DHE) dye. Slides were incubated in 10 mM DHE for 10 minutes at 37°C, washed with 1X PBS, and nuclei were stained with DAPI. Images were captured using a Nikon Ti2 ECLIPSE confocal microscope at 40X magnification, 1.4 aperture, and further analyzed using the Nikon NIS Elements software. DHE intensity was quantified using 3 different mBEDO samples, and 5 ROIs were averaged to obtain mean fluorescence intensity in ImageJ software.

### 2.14 BMDM co-culture with organoids

To generate mouse BMDMs, bone marrow was flushed from the mouse tibia and femur with a 27G ½ inch syringe needle and 1XPBS supplemented with 1% BSA as described previously(Symington et al., 2015). Isolated marrow was passed through 70 µm cell strainer and mashed with the syringe plunger. 1XRBC lysis buffer (420301, BioLegend) was used to remove red blood cells. 1-2×10^6^ cells were plated in 10 cm bacterial Petri dish in DMEM/F12 media supplemented with 20% FBS, 50 ng/mL mM-CSF (315-02, Peprotech) and penicillin/streptomycin antibiotics. After 7 days of culture, cells were collected and BMDMs differentiation rate examined with flow cytometry surface staining by the enrichment of CD11b+ F4/80+ double positive cell population (101205, BioLegend for CD11b-FITC; 02922-80-25, biogems for F4/80-APC). Co-culture experiment setup was adopted from Noel, Baetz, et al.,(Noel et al., 2017)with modifications. 0.2-2×10^6^ BMDMs were plated in differentiation bladder organoids medium supplemented with 50ng/mL of mM-CSF on inverted 0.4 um PET transwell inserts (3470, Costar) in 6-well plates, then incubated for 2 hours at 37C. The PET inserts with attached BMDMs were placed back to a normal position. 100,000 differentiated bladder organoids were added on top of the PET inserts to grow as a monolayer in 24-well plates in differentiation medium supplemented with 50ng/mL of mM-CSF. Cells were co-cultured overnight prior to downstream analysis. To perform a whole-mount immunofluorescent staining, inserts were rinsed with 1XPBS twice at RT, then fixed with 4% of methanol-free PFA at RT for 5 min, followed by two washes with 1XPBS. After fixation, the membrane was blocked with 1%BSA plus 0.1% Triton X-100 for 1 hour at RT. After blocking, the primary antibodies were added for overnight incubation at 4C (ab53121, Abcam for Cytokeratin 5 1:200; ab6640, Abcam for F4/80 1:300). Prior to adding fluorescently labelled secondary antibodies, the membrane was washed 3 times with 1XPBS. The membrane was incubated for 1 hour at RT with secondary antibodies, followed by three washes with 1XPBS and mounting the whole membrane in ProLong Gold antifade with DAPI mounting solution (P36931, Thermo Fisher Scientific). Images were captured using a Nikon Ti2 ECLIPSE confocal microscope at 40X magnification, 1.4 aperture, and further analyzed using the Nikon NIS Elements software.

### 2.15 Metabolite supplementation of mBEDOs

Aged mBEDOs were supplemented with 17.5 mM D-Mannose and 1mM nicotinamide (NAM) in Advanced DMEM, 10% FBS, Pen/Strep and GlutaMAX. Organoids were harvested at 24 hours post-treatment with each metabolite. Individual Matrigel tabs were collected at the 24-hour timepoint, Matrigel was removed and organoids were either frozen in OCT to perform histological analysis and staining or pelleted and stored in TriZOL for RNA extraction, cDNA synthesis, and qRT-PCR.

### 2.16 Untargeted metabolomics of mBEDOs

Organoids were extracted out of Matrigel and frozen pellets were stored at −80°C. Organoids were homogenized using needle sonication in a 50:50 methanol/water mixture. Subsequently, three volumes of methanol/acetonitrile (50:50, v/v) were added to the samples. The samples were vortexed for 5 minutes and then stored at 20°C for 10 minutes. Afterward, they were centrifuged at 15,000 rpm for 10 minutes at 4°C. The supernatants were evaporated to dryness using a GeneVac EZ-2 Plus SpeedVac (SP Scientific). The dried samples were reconstituted in 100 µL of methanol/water (50:50, v/v). A pooled quality control (QC) sample was prepared by mixing 20 µL of each supernatant and dividing it into aliquots.

Separation was performed using a Thermo Scientific Vanquish Horizon UHPLC system. A Waters ACQUITY HSS T3 column (1.8 μm, 2.1 mm × 150 mm) was used for reversed-phase separation, while a Waters ACQUITY BEH Amide column (1.7 μm, 2.1 mm × 150 mm) was employed for HILIC separation. For reversed-phase separation, solvent A consisted of 0.1% formic acid in water, and solvent B consisted of 0.1% formic acid in methanol. For HILIC separation, solvent A was prepared with 0.1% formic acid, 10 mM ammonium formate, 90% acetonitrile, and 10% water, while solvent B contained 0.1% formic acid, 10 mM ammonium formate, 50% acetonitrile, and 50% water. Both columns were operated at 50 °C with a flow rate of 300 μL/min and an injection volume of 2 μL. Data were collected using a Thermo Scientific Orbitrap IQ-X Tribrid Mass Spectrometer equipped with an H-ESI source. A spray voltage of 3500 V was applied for positive ion mode during reversed-phase separation, and 2500 V was used for negative ion mode during HILIC separation. The vaporizer temperature and ion transfer tube were maintained at 300 °C.

The dataset was processed using Compound Discoverer (v3.3, Thermo Fisher Scientific) for peak detection, integration, and identification. Metabolites were identified using mzCloud, the NIST 2020 HRMS library, the in-house retention time-based library, and the HMDB mass list. Peak areas were log2 transformed and normalized using the median IQR method per method. Differential metabolites were identified with p-values < 0.05.

### 2.17 Quantification and statistical analysis

An “n” of 3 was used for all fluorescence intensity measurements and an “n” of 6 was used for qRT-PCR analysis. The “n” are biological replicates representing mBEDOs derived from a single mouse, grown in a single well of a 6-well plate. All measured values were analyzed using GraphPad Prism versions 9.0.1 to 10.2.2 (GraphPad Software, La Jolla, CA, USA). Two-tailed unpaired t-tests were used when data approximated normal distribution. “n” for each experiment and statistical test used are indicated in figure legends. Data were expressed as ± Standard Error of the Mean (SEM).

## 3 Results

### 3.1 mBEDOs recapitulate key features of *in vivo* urothelium

We generated organoids using whole bladders from young (2-3 months) and aged (15–18 months) female C57BL/6J mice (**Figure 1A**). Briefly, bladder tissue was dissociated in a cocktail of Collagenase II, Elastase, and Y-27632. Single cells were harvested by centrifugation and prior cocktail supernatant was replaced with TrypLE for further digestion. After gentle centrifugation, single cells were resuspended in Advanced DMEM and a total of 20,000 cells were suspended in 50 µL Matrigel tab and plated in 6-well non-treated cell culture plates (4-5 tabs per well). Organoids were generated through a 3-stage protocol (**Figure 1A**). In the first stage, cells were maintained for seven days in a proliferation medium enriched with growth factors including mouse EGF, Noggin, and R-Spondin. During this time, we observed initial increases in cell number followed by the formation of spheroids. On day 7, organoids were transitioned to an intermediate medium containing WNT3A and IL-6, which promoted cellular maturation and halted proliferation, leading to an increase in spheroid size. Finally on day 14, organoids were moved to differentiation medium supplemented with FBS, FGF-7, and FGF-10 to support terminal differentiation. By day 21, organoids developed into fully differentiated, mature, multi-layered structures with a central lumen, consistent with urothelial tissue organization *in vivo* (**Figure 1A**). At the end of the proliferation stage, cells were subcultured to facilitate scalability and technical replication. New tabs were made to maintain a 20,000 cells/tab density throughout the rest of the organoid development protocol. In addition, some cells were gently extracted from the Matrigel tab for cryopreservation to support longitudinal studies.

**Figure 1.**
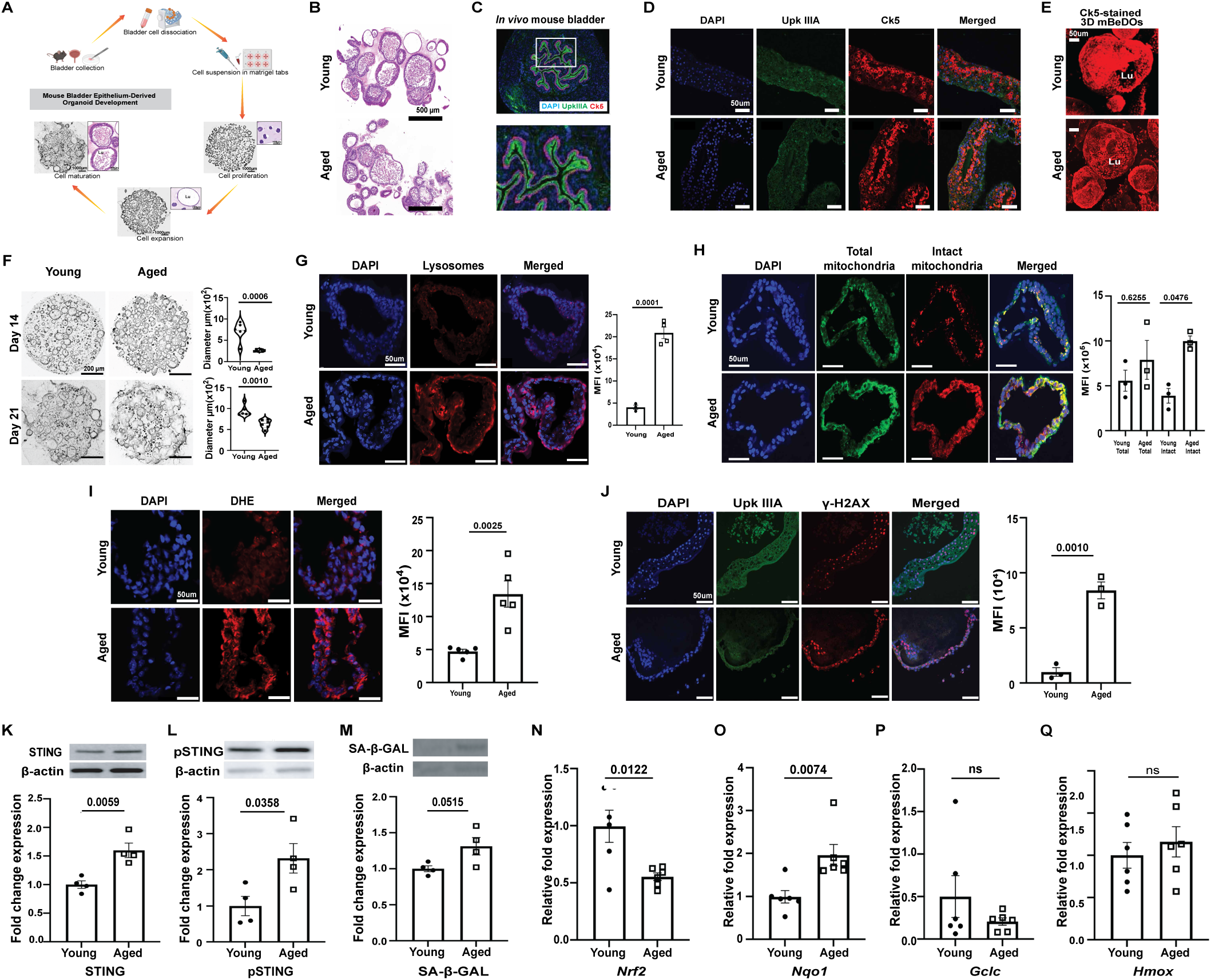
**mBEDOs recapitulate several key features of *in vivo* urothelium**. (A) Schematic diagram of mBEDO generation. (B) Hematoxylin and Eosin staining of wild-type young and aged mBEDOs show 3D cellular architecture and makeup of mBEDOs (n=3/group), scale bar = 500 µm. (C) Expression of UPKIIIA and CK5 in *in vivo* bladder urothelium (D) UPKIIIA and CK5 staining following mBEDO fixation revealing UPKIIIA and CK5 expression in young and aged mBEDOs, scale bar = 50µm. (E) CK5 staining of 3D mBEDOs revealing a robust expression in all cells at homeostasis, scale bar = 50µm. (F) Brightfield images of mBEDOs in Matrigel displaying differences in organoid diameters at days 14 and 21 mBEDO development, scale bar = 200µm. (G) LysoTracker staining of fresh-frozen mBEDOs revealing lysosomal accumulation in aged mBEDOs, scale bar = 50µm. (H) Mitochondrial staining using Mito Tracker Green FM and MitoTracker Orange CMTM ROS showing total mitochondrial mass is higher in aged mBEDOs, scale bar = 50µm. (I) DHE staining revealing higher levels of superoxide radicals in aged mBEDOs compared to young, scale bar = 25µm. (J) UPKIIIA and γH2AX staining in young and aged mBEDOs (Data presented as mean ± SEM, n = 3 each group). (K) Western Blot quantitation of STING in young and aged mBEDOs (n = 4 each group). (L) Western Blot quantitation of pSTING in young and aged mBEDOs (n = 4 each group). (M) Western Blot quantitation of SA-β-gal in young and aged mBEDOs (n = 4 each group). Real-time quantitative PCR analysis of (N) *Nrf2*, (O) *Nqo1*, (P) *Gclc*, and (Q) *Hmox* in young and aged mBEDOs (Data presented as mean ± SEM, n = 6 for *Nrf2*, *Nqo*, *Gclc* and *Hmox*).

Hematoxylin and Eosin (H&E) staining showed that differentiated mBEDOs developed a multilayer architecture with a central lumen, similar to bladder urothelium (**Figure 1B-C**). Immunofluorescence staining demonstrated the presence of both basal (Cytokeratin 5; CK5) and apical (Uroplakin IIIA; UPKIIIA) cell markers (**Figure 1D**), while three-dimensional imaging further confirmed the robust expression of CK5 in both young and aged mBEDOs (**Figure 1E**). We evaluated growth dynamics of mBEDOs using organoid diameter, as a measure of growth, during the proliferation (Day 14) and differentiation (Day 21) stages and found that aged mBEDOs exhibited significantly smaller diameters than young counterparts at both time points (Day 14: *p* = 0.0006; Day 21: *p* = 0.0010) (**Figure 1F**). This observation reflects prior findings of reduced growth capacity in aged tissues(Carlson & Conboy, 2007) and further adds a feature of *in vivo* aged urothelium recapitulated in mBEDOs.

Our previous work has demonstrated that the aged urothelium *in vivo* exhibits cellular and molecular alterations when compared to young such as dysfunctional lysosomal and mitochondrial compartments, increased reactive oxygen species (ROS) generation, decreased antioxidative response, and senescence associated phenotype(Joshi et al., 2024; Ligon et al., 2020). We determined whether these features were reflected in mBEDOs. We stained for lysosomes (using LysoTracker) from fresh frozen sections. 5 mean fluorescence intensity (MFI) readings per organoid from a single well and single replicate (each well represents a biological replicate which corresponds to mBEDOs derived from a single mouse). We observed a significant five-fold increased MFI in aged compared to young mBEDOs signifying an increase in lysosomal abundance (p = 0.001) (**Figure 1G**). A separate staining for total mitochondria (Green FM MitoTracker – membrane potential-independent) and intact mitochondria (Orange CMTM ROS – membrane potential-dependent) revealed no significant change in the total mitochondria between young and aged mBEDOs; however, a 40% significantly increased MFI for intact mitochondria populations was observed in the latter (p = 0.0476) (**Figure 1H**). Using dihydroethidium (DHE) stain, we observed a 2.8-fold increase in reactive oxygen species (ROS) in aged mBEDOs compared to young (p = 0.025) (**Figure 1I**). Downstream consequences of chronic oxidative stress include the accumulation of DNA damage and cellular senescence, both well-established hallmarks of aging observed in multiple tissues, including the aged bladder urothelium(Joshi et al., 2024; Maldonado et al., 2023; Schumacher et al., 2021). Using γH2AX – a marker of DNA double-strand breaks, we assessed the level of DNA damage in fixed mBEDO samples. Significantly higher MFI for γH2AX-positive nuclei was observed in aged compared to young mBEDOs (p = 0.001) (**Figure 1J**). Chronic DNA damage and oxidative stress can also activate the endoplasmic reticulum (ER)-resident protein STING (stimulator of interferon genes), a driver of innate immune signaling and senescence^19^. Aged mBEDOs displayed significantly higher levels of both total and phosphorylated STING compared to young organoids **(Figure 1 K-L**). In aging, activated STING can influence the regulation of Senescence-associated Secretory Proteins (SASP) (Zheng et al., 2023; T. Li & Chen, 2018). Senescent cells typically accumulate senescence-associated β-galactosidase (SA-β-Gal) – a widely used marker that also correlates with increased lysosomal content (Dimri et al., 1995; Debacq-Chainiaux et al., 2009; Lee et al., 2006). Using western blot, we found SA-β-Gal was significantly elevated in aged mBEDOs compared to their young counterparts by 30% (p = 0.0389) (**Figure 1M),** consistent with increased senescence reported in the aged urothelium (Joshi et al., 2024).

We previously reported that while ROS accumulates in the aged bladder, the oxidative stress is not adequately counteracted due to incomplete activation of the NRF2 antioxidant program (Joshi et al., 2024). NRF2 – a key regulator of cellular redox homeostasis, is typically suppressed under homeostatic conditions in urothelial cells but becomes activated in response to increased ROS levels (Joshi et al., 2021). Using quantitative Real-Time Polymerase Chain Reaction (qRT-PCR), we assessed gene expressions of nuclear factor erythroid 2-related factor 2 (NRF2) and its canonical target genes in young and aged mBEDOs. Aged mBEDOs showed significantly lower levels of *Nrf2* compared to young by 44% (p = 0.0122) (**Figure 1N),** indicating a dampened antioxidant response in aged mBEDOs. *Nqo1* was increased in aged compared to young mBEDOs (p = 0.0074) and no changes in *Gclc* and *Hmox* expression were noted (**Figure 1O, P, Q**).

These findings mirror our *in vivo* results and support the conclusion that aged urothelial cells experience elevated oxidative stress that is not sufficiently mitigated by a robust NRF2-mediated antioxidant response. Together, these findings verify that mBEDOs recapitulate multiple molecular and structural hallmarks of urothelial aging, establishing them as a robust and scalable *ex vivo* model for studying epithelial aging.

### 3.2 mBEDO platform demonstrates receptivity to UPEC infection, shows interaction with macrophages, and suitability for gene knockout organoid development

Uropathogenic *Escherichia coli* (UPEC) is the most common bacterial infection that accounts for 80-90% of UTI cases in the world (Whelan et al., 2023), with increasing frequency and severity with age. We previously demonstrated that aged female mice, exhibit elevated susceptibility to recurrent UTIs, which has been linked to altered urothelial barrier function, inflammation, and immune dysregulation (Joshi et al., 2024). UPEC initiates infection by adhering to and invading Uroplakin-positive cells of the urothelium (Liu et al., 2015). We next examined whether mBEDOs could be infected with UPEC. We employed 2D monolayers to allow and maximize direct bacterial contact with Uroplakin-positive cells (**Figure 2A**). Briefly, differentiated 3D organoids were enzymatically dissociated into single cells using TrypLE and seeded glass-bottom plates, pre-coated with thin layer of Matrigel for four days. Monolayers were infected with GFP-expressing UPEC at an MOI of 1:100 for 1- or 3-hours, and at each time point they were treated with gentamicin to eliminate extracellular bacterial growth. This also ensured that infections were from the bacteria that infiltrated the Uroplakin-positive cells. Confocal imaging revealed GFP-labelled UPEC (green) infected and localized to Uroplakin-positive cells (orange) in both young and aged 2D mBEDOs (**Figure 2B**). Aged mBEDOs however, show significantly higher MFI compared to young at 3 hpi (p = 0.005), indicating increased bacterial accumulation over time (**Figure 2C**). Bacterial load quantification via colony forming units further demonstrated significantly higher intracellular bacterial population in aged mBEDOs (**Figure 2D**).

**Figure 2.**
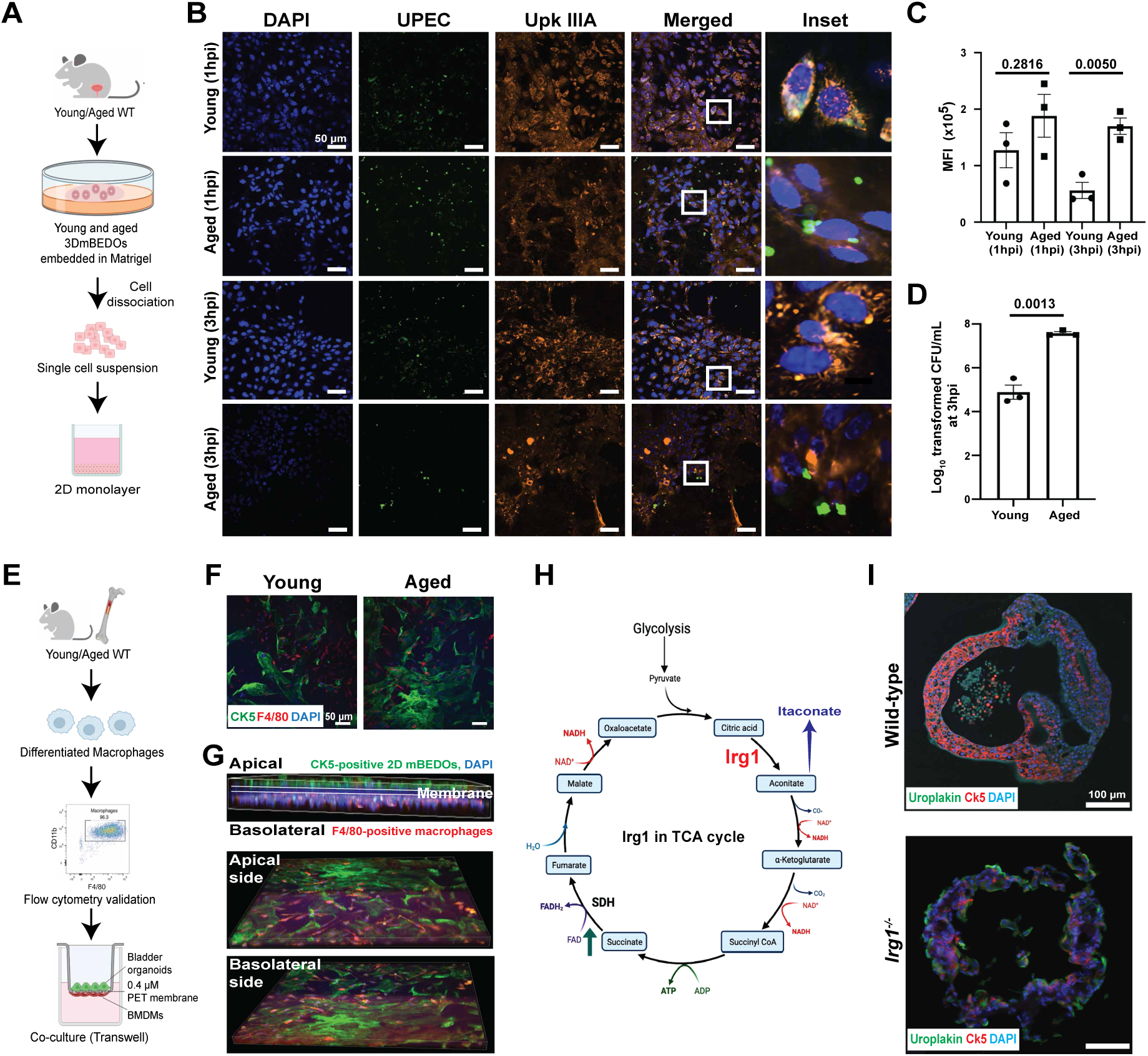
mBEDOs model UPEC infection, epithelial-immune cell interactions, and genetic knockouts. (A) Schematic representing UPEC infection of 2D mBEDO monolayers. (B) Representative images of IF localization of E. coli (green) forming bacterial clusters in young and aged mBEDOs. UPKIIIA (yellow) marks cell membrane and DAPI (blue) stains nuclei. (C) Quantitation of number of bacteria at 1 and 3 hpi in young and aged mBEDOs (Data presented as mean ± SEM, n = 3 in each group, p values by two-tailed unpaired t test). (n=3/group), scale bar = 50 µm. (D) Bacterial colony forming units (CFUs) at 3 hpi in young and aged 2D mBEDO monolayers (Data presented as mean ± SEM, n = 3 in each group, p values by two-tailed unpaired t test). (E) Schematic representing BMDM isolation and co-culture with UPEC via transwell inserts. (FF) Representative image of BMDMs on the basolateral side co-localizing with urothelial monolayers on the apical side of transwell inserts. (GG) Representative images of IF localization of BMDMs (red) with young and aged mBEDOs. CK5 (green) marks cell membrane and DAPI (blue) stains nuclei, scale bar = 50 µm. (H) TCA cycle dynamics at homeostasis, *Irg1* activation with aging and subsequent production of itaconate. (I) Representative images of wild-type young and *Irg1^-/-^* mBEDOs showing CK5 (red) and UPKIIIA (green). DAPI (blue) stains nuclei, scale bar = 100 µm.

Given the complex immune landscape in the bladder particularly its network of resident macrophages positioned beneath the urothelium (A. S. Wang et al., 2022), we sought to examine epithelial–immune cell interactions using the mBEDO platform. Macrophages are known to have primary function in immune homeostasis, which declines with age (A. S. Wang et al., 2022). Using transwell inserts, we co-cultured young and aged 2D monolayers with bone marrow–derived macrophages (BMDMs) from young and aged mice (**Figure 2E**). This system enables interaction between urothelial cells (CK5-positive) on the apical compartment with the macrophages (F4/80-positive) on the basolateral compartment. Confocal imaging revealed colocalization of macrophages with urothelial cells (**Figure 2F**). These interactions were observed in young and aged mBEDO-assembloids (**Figure 2G)**, demonstrating the utility of this platform for studying epithelial-immune cell dynamics.

Aging is associated with significant metabolic reprogramming, including alterations in the tricarboxylic acid (TCA) cycle (Borkum, 2023; Kurhaluk, 2024). One key response to this dysregulation is the induction of *Irg1* (also known as *Acod1*)(Wu et al., 2022), which catalyzes the production of the immunometabolite, itaconate from aconitate (R. Li et al., 2020) (**Figure 2H**). Elevated itaconate levels have been linked to chronic inflammation and tissue remodeling, but its role in urothelial aging remains unknown. Whole bladder RNAseq of young and aged bladders (Ligon et al., 2020) revealed that *Irg1* is upregulated with age, suggesting a contribution to age-associated metabolic shifts (**Figure S1**); however, its specific role in urothelial aging remains unknown. To test whether the mBEDO platform could be extended to interrogate gene function in aging, we generated mBEDO from 2-3 month old female *Irg1⁻/⁻*mice. Immunofluorescence staining confirmed the presence of key urothelial markers, with expression of basal marker CK5 and apical marker UPKIIIA present in both *Irg1^⁻/⁻^* and young wild type mBEDOs (**Figure 2I**). Our results demonstrate the potential of the mBEDO platform for interrogating underpinnings of genetic modification on urothelial dynamics.

### 3.3 Distinct and significant metabolic signatures in young, aged and *Irg1^⁻/⁻^* mBEDOs

Metabolic dysregulation is a hallmark of aging (Zhang et al., 2021). Here, we sought to determine whether the mBEDO platform could be leveraged to define metabolic signatures associated with aging. We performed untargeted metabolomics on young and aged mBEDOs to identify pathways perturbed with age and to evaluate candidate metabolites for therapeutic supplementation. mBEDOs from young and aged bladders were isolated from all Matrigel tabs present in a single well. Each well represents a biological replicate with organoids derived from a single mouse. Principal component analysis (PCA) of metabolite abundances revealed clear and distinct global metabolic profiles between young and aged mBEDO samples (**Figure 3A**) with 20 metabolites significantly decreased and 21 increased in aged compared to young mBEDOs (**Figure 3B**). In aged mBEDOs, mannose and ketobutyric acid were reduced while adenosine, guanine, hypoxanthine, inosine, and choline were significantly increased (**Figure 3B**). We additionally performed pathway enrichment analysis to identify overrepresented pathways that may unravel the cellular and molecular differences between young and aged mBEDOs. Pathways were ranked by enrichment significance and impact score, with node size and color reflecting the degree of metabolic disruption. Our analysis revealed significant enrichment of glycerophospholipid metabolism, purine metabolism, and amino acid metabolism, specifically that involving glycine, serine, and threonine (p < 0.05) (**Figure 3C**).

**Figure 3.**
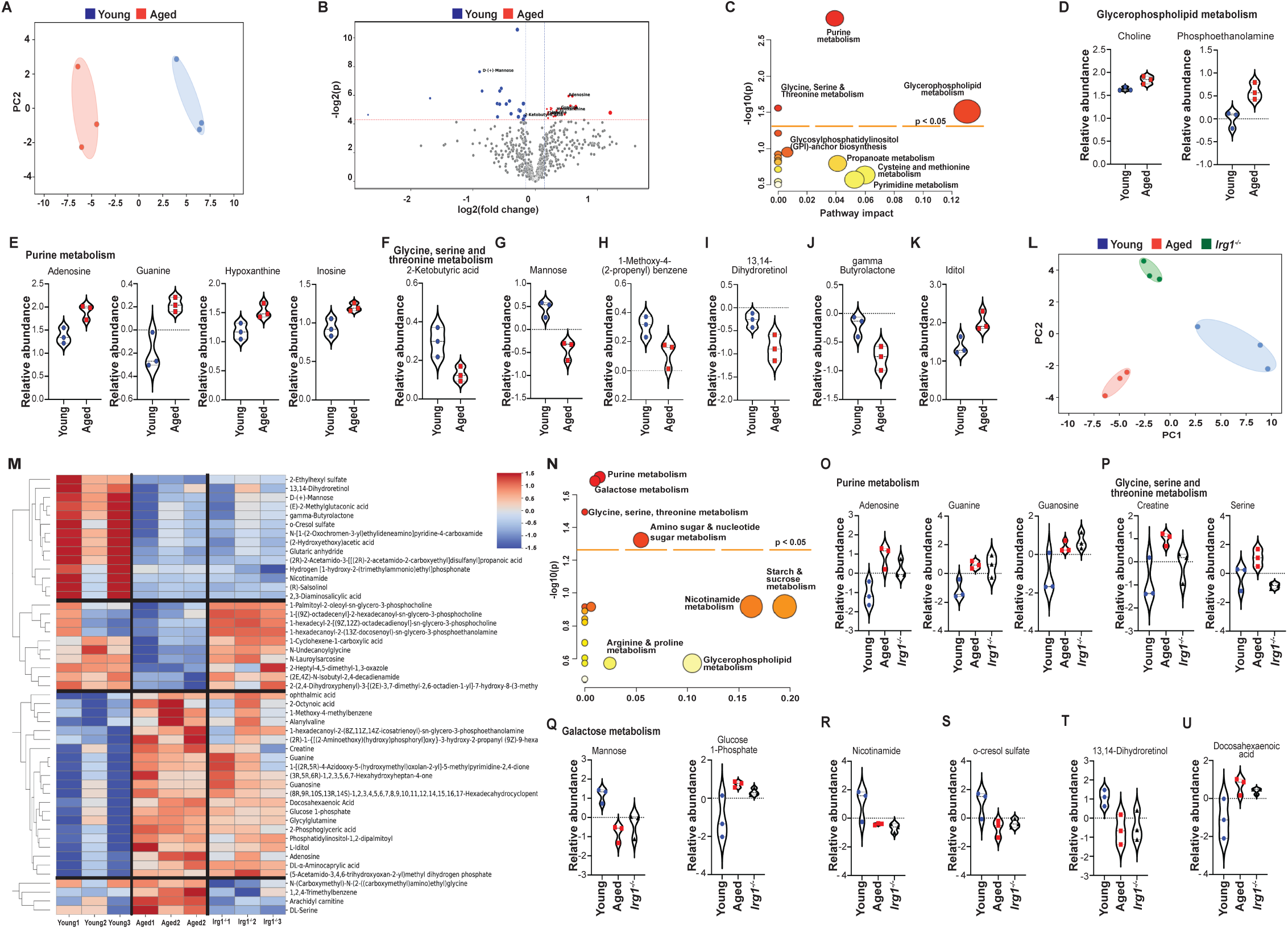
Young, Aged and *Irg1^-/-^* mBEDOs, have stark differences in their metabolic profiles. (A) PCA plot where the red dots denote the aged mBEDO samples and the blue dots denote those in the young group. The ellipses illustrate the range of sample variability per group. (B) Volcano plot displaying overall metabolites that were differentially and significantly (p<0.05) altered between young and aged samples. Red indicates higher relative abundance and blue indicates lower relative abundance for each metabolite. (C) Pathway impact map displaying metabolite pathways that were significantly represented in the metabolomics data. Size of the circles directly corelates to the number of metabolites representing a pathway. (D) Relative abundances of metabolites from the glycerophospholipid pathway. Red color indicates aged while blue indicates young samples. (E) Relative abundances of metabolites from the purine metabolism pathway. Red color indicates aged while blue indicates young samples. (F) Relative abundance of a metabolite mapped to glycine, serine, threonine metabolism. Red color indicates aged while blue indicates young samples. (G) Relative abundance of Mannose in young (blue) and aged (red) samples. Red color indicates aged while blue indicates young samples. (H) Relative abundance of 1-Methoxy-4-(2-propenyl) benzene in young (blue) and aged (red) samples. Red color indicates aged while blue indicates young samples. (I) Relative abundance of 13,14-Dihydroretinol in young (blue) and aged (red) samples. Red color indicates aged while blue indicates young samples. (J) Relative abundance of gamma Butyrolactone in young (blue) and aged (red) samples. Red color indicates aged while blue indicates young samples. (K) Relative abundance of Iditol in young (blue) and aged (red) samples. Red color indicates aged while blue indicates young samples. (L) PCA plot where the red dots denote the aged mBEDO samples, blue dots denote those in the young group, and green are the *Irg1^-/-^* mBEDOs. The ellipses illustrate the range of sample variability per group. (M) Heatmap displaying overall metabolites that were differentially and significantly (p<0.05) altered between young, aged, and *Irg1^-/-^* samples. Red indicates higher relative abundance and blue indicates lower relative abundance for each metabolite. (N) Pathway impact map displaying metabolite pathways that were significantly represented in the metabolomics data. Size of the circles indicate more metabolites representing a pathway. (O) Relative abundances of metabolites from the purine metabolism pathway. Red color indicates aged, blue indicates young and green indicates *Irg1^-/-^* samples. (P) Relative abundance of a metabolite mapped to glycine, serine, threonine metabolism. Red color indicates aged, blue indicates young and green indicates *Irg1^-/-^* samples. (Q) Relative abundance of a metabolite mapped to galactose metabolism. Red color indicates aged, blue indicates young and green indicates *Irg1^-/-^* samples. (R) Relative abundance of NAM. Red color indicates aged, blue indicates young and green indicates *Irg1^-/-^* samples. (S) Relative abundance of o-cresol sulfate. Red color indicates aged, blue indicates young and green indicates *Irg1^-/-^* samples. (T) Relative abundance of 13,14-Dihydroretinol. Red color indicates aged, blue indicates young and green indicates *Irg1^-/-^* samples. (U) Relative abundance of Docosahexaenoic acid. Red color indicates aged, blue indicates young and green indicates *Irg1^-/-^*samples. Data presented as mean ± SEM, n = 3 each group.

We further examined these pathways to identify significant changes in specific individual metabolite abundances. In the glycerophospholipid metabolism pathway, 1-hexadecanoyl-2-(8Z,11Z,14Z-icosatrienoyl)-sn-glycero-3-phosphoethanolamine (PE(16:0/20:4)) and choline were elevated in aged mBEDOs when compared to young (**Figure 3D**). In the purine metabolism pathway, levels of inosine, hypoxanthine, guanine, and adenosine were also significantly higher in aged mBEDOs (**Figure 3E**). However, 2-ketobutyric acid, a metabolite mapped to glycine, serine, and threonine metabolism, was reduced in aged mBEDOs (**Figure 3F**). We also identified several biologically relevant metabolites that were present in the KEGG database but not confidently assigned to a specific pathway in MetaboAnalyst. Mannose (D or L) (**Figure 3G**), 1-methoxy-4-(2-propenyl) benzene (**Figure 3H**), dihydroretinol (**Figure 3I**), and gamma-butyrolactone (**Figure 3J**) were all significantly decreased in aged mBEDOs. Additionally, Iditol (sorbitol) was elevated in aged mBEDOs (**Figure 3K**), possibly reflective of the increased ROS levels. Together, our results reveal broad and specific shifts in metabolic signatures between young and aged mBEDOs, highlighting altered redox, nucleotide, lipid, and carbohydrate metabolism as potential drivers or consequences of urothelial aging.

To assess the influence of genetic perturbation of IRG1/itaconate on metabolic aging, we compared the metabolic profiles of young, aged, and *Irg1^⁻/⁻^* mBEDOs. PCA plots revealed three separate and distinct clusters representing each group (**Figure 3L**) with 21 metabolites that were significantly increased and 15 metabolites that were decreased in aged and *Irg1^-/-^* mBEDOs compared to young (**Figure 3M**). In addition, we observed 10 metabolites that were significantly increased in the young and *Irg1^-/-^* mBEDOs but were decreased in aged mBEDOs. Further, we found 4 metabolites that were lower in abundance in both young and *Irg1^-/-^* mBEDOs but increased in the aged mBEDOs (**Figure 3M**).

Next, we conducted pathway enrichment analysis on the significantly altered metabolites. We identified enrichment in purine, galactose, glycine–serine–threonine, and nucleotide sugar metabolism pathways (**Figure 3N)**. Upon closer inspection of individual metabolites, specific overlaps between aged and *Irg1⁻/⁻* mBEDOs were evident. In purine metabolism, adenosine, guanine, and guanosine were significantly elevated in both aged and *Irg1⁻/⁻* mBEDOs when compared to young (**Figure 3O**). In amino acid metabolism, D/L-serine and creatine were increased in aged but not in *Irg1⁻/⁻*mBEDOs, suggesting partial divergence in metabolic signatures (**Figure 3P**). Further in galactose metabolism, we found lower mannose abundance in aged and *Irg1^-/-^*mBEDOs when compared to young (**Figure 3Q**). However, in galactose metabolism, glucose-1-phosphate (**Figure 3Q**) was higher in aged and *Irg1^-/-^* mBEDOs compared to the young. We also examined additional biologically relevant metabolites (identified via KEGG) that were significantly altered but not confidently assigned to a single pathway. Among these, nicotinamide, o-cresol sulfate, and 13,14-Dihydroretinol were reduced in aged and *Irg1⁻/⁻* mBEDOs (**Figure 3R-T**), while docosahexaenoic acid (DHA) was elevated in both aged and *Irg1⁻/⁻* mBEDOs relative to young (**Figure 3U**).

Together, these findings demonstrate that IRG1 deficiency in young mBEDOs recapitulates many of the metabolic alterations observed in mBEDOs generated from aged bladders, particularly in purine and galactose metabolism. These data suggest a role for IRG1 in maintaining urothelial metabolic homeostasis and that IRG1 loss-of-function may act as a metabolic accelerator of aging phenotypes in the bladder.

### 3.4 Metabolite supplementation alleviates oxidative stress and DNA damage in aged mBEDOs

To explore the utility of the mBEDO platform for therapeutic development and assessment, we supplemented aged mBEDOs with NAM and D-mannose, two metabolites identified as significantly depleted in our untargeted metabolomics screen. Notably, D-mannose has been previously reported by our group to exert senotherapeutic effects in the aged female bladder *in vivo*, while NAM and its precursors have been widely implicated in promoting health span and lifespan extension across multiple model systems(Imai & Guarente, 2014; Verdin, 2015; Anderson et al., 2003). We hypothesized that supplementation of depleted metabolites would improve the aging phenotypes observed in mBEDOs. At the end of the differentiation stage, aged mBEDOs were cultured in medium containing either metabolite followed by harvesting after 24 hours. Samples were processed for histological analysis, live imaging, and gene expression profiling. We exposed young mBEDOs as well to further assess the response to above-normal levels of metabolites and found no adverse effects. We examined lysosomal abundance, a feature known to increase with age, and neither D-mannose nor NAM showed significant changes either in young and aged mBEDOs when compared to untreated mBEDOs (**Figure 4A-B).** We next assessed mitochondrial population by staining for intact mitochondria (membrane potential-dependent; red) and total mitochondrial (membrane potential-independent; green) contents (**Figure 4C**). Aged mBEDOs show significant reduction in intact mitochondria when treated with both NAM and D-mannose treatments. NAM and D-mannose treatments resulted in a fourfold (p = 0.0186) and threefold (p = 0.0103) decrease, respectively. Similarly, Total mitochondria were significantly reduced following treatment with NAM and D-mannose (p < 0.0001). These observations suggest a shift in mitochondrial dynamics with supplementation. Young mBEDOs exhibited lower total mitochondrial population in both treatment settings (NAM, *p* = 0.0011; D-mannose, *p* = 0.0004) when compared to the untreated group (**Figure 4D**). We also assessed ROS levels by DHE staining. Both NAM and D-mannose treatments significantly reduced ROS levels in aged mBEDOs when compared to untreated groups (NAM, *p* = 0.0026; D-mannose, *p* = 0.0183) **(Figure 4 E & F**), indicating an improvement in redox homeostasis. This reduction in oxidative stress is consistent with the observed reduction in mitochondrial abundance in both treatment groups. Finally, we evaluated DNA damage via γH2AX staining. Both treatments showed significant reduction in DNA damage marker intensity in aged mBEDOs (D-mannose and NAM *p* < 0.0001) (**Figure 4 G & H**), suggesting partial decline in aging-associated DNA damage. Together, our results demonstrate that mBEDOs provide a scalable platform for performing metabolite discovery and supplementation to improve specific aging phenotypes.

**Figure 4.**
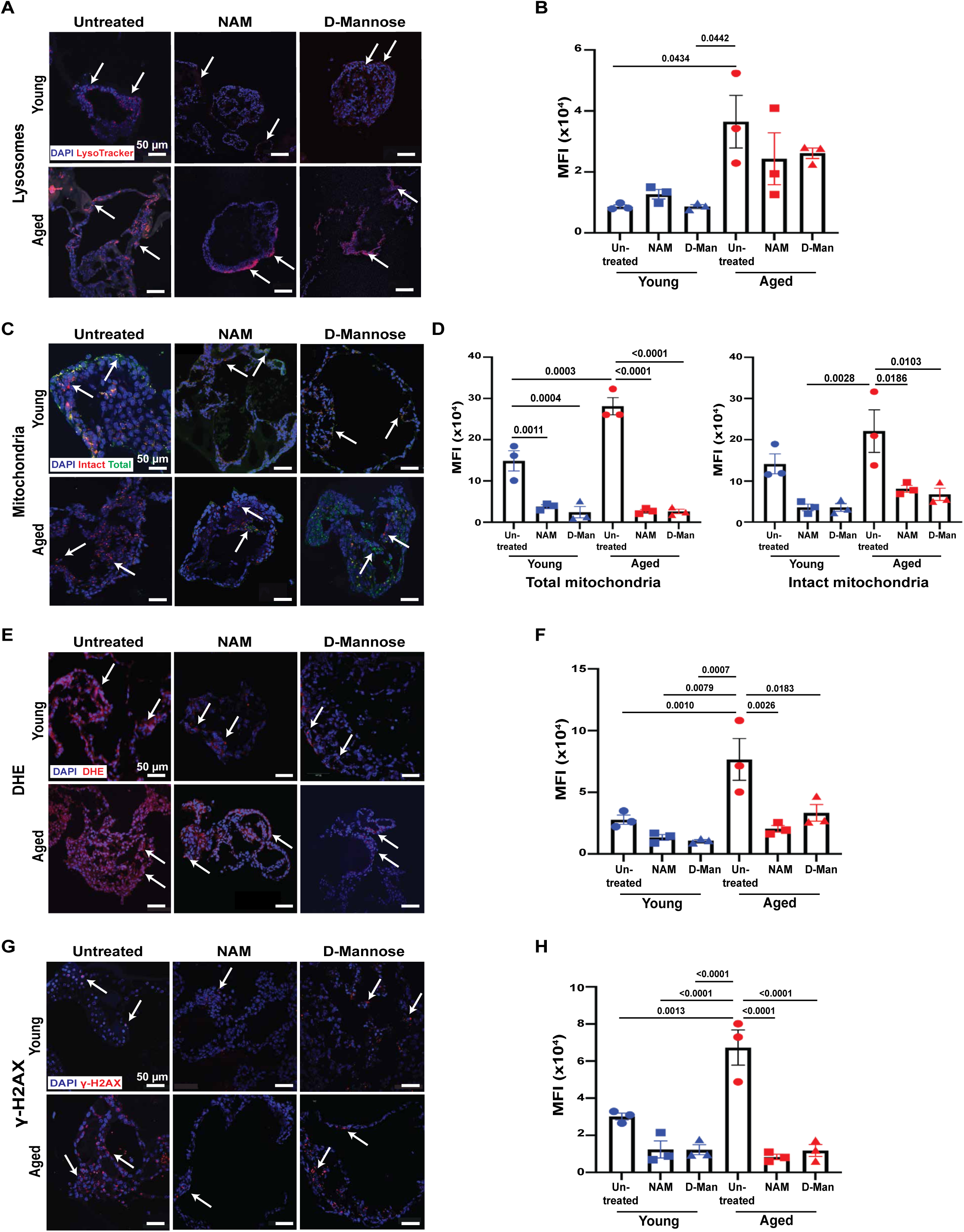
Metabolite supplementation alleviates oxidative stress in aged mBEDOs. (A) LysoTracker staining of fresh-frozen mBEDOs revealing lysosomal accumulation (Red) in young and aged untreated and treated with NAM and D-Mannose, scale bar = 50µm. (B) Quantitation of Lysotracker staining in young and aged untreated mBEDOs and those treated with NAM and D-Mannose (Data presented as mean ± SEM, n = 3 each group) (C) Mitochondrial staining using Mito Tracker Green FM and MitoTracker Orange CMTM ROS showing total mitochondrial (Green) and intact mitochondria (Red), in young and aged untreated and treated with NAM and D-Mannose mBEDOs, scale bar = 50µm. (D) Quantitation data for mitochondrial staining in all treated and untreated groups (Data presented as mean ± SEM, n = 3 each group). (E) DHE staining showing reactive oxygen species (Red), in young and aged untreated and treated with NAM and D-Mannose mBEDOs, scale bar = 50µm. (F) Quantitation data for mitochondrial staining in all treated and untreated groups (Data presented as mean ± SEM, n = 3 each group). (G) **γ**-H2AX (Red) staining showing double-stranded DNA breaks in young and aged untreated and treated with NAM and D-Mannose mBEDOs, scale bar = 50µm. (H) Quantitation data for mitochondrial staining in all treated and untreated groups (Data presented as mean ± SEM, n = 3 each group). DAPI stains all nuclei blue.

## 4 Discussion

Advancements in 3D organoid cultures have improved our understanding of lower urinary tract biology. These reports have enabled a wide range of studies ranging from urothelial responses to UPEC infection to the recapitulation of the bladder cancer tumor microenvironment; Mullenders et al., 2019). In this study, we present a practical, scalable, and reproducible platform of generating organoids from mouse whole bladders that facilitates investigations geared towards elucidating aging-related urothelial mechanisms. Multiple characteristics of the *in vivo* mouse aged bladders are retained or reflected by the aged mBEDOs such as reduced growth, increased lysosomal and mitochondrial accumulation, elevated oxidative stress, increased DNA damage and activation of innate immune response, insufficient antioxidative response, and increased senescence. Our 3-stage mBEDO protocol consisting of proliferation, cell expansion, and cell differentiation, demonstrates robust production of differentiated, multilayer, and central lumen-containing organoids providing an opportunity to study urothelial-intrinsic phenotypes. Our model adds to the growing body of *ex vivo* organoid models (Smith et al., 2006; Mullenders et al., 2019; E. Kim et al., 2020; Azar et al., 2021; Torrens-Mas et al., 2021; Sato et al., 2009; Walz et al., 2023) with the ability to not only gain insights into tissue-specific aging but also provide metabolic insights leading to therapeutic interventions specifically targeting urothelial aging hallmarks.

Several studies have utilized organoid models to explore the interplay between immune cells and epithelial cells. Intestinal organoids have been used to investigate how immune cells such as T cells and macrophages interact with the epithelial layer in response to infections or inflammatory stimuli (Recaldin et al., 2024; Múnera et al., 2023). In addition, gastric organoids have been employed to study infections caused by *Helicobacter pylori*, while lung organoids help to model respiratory infections, such as those caused by *Salmonella enterica* or the respiratory syncytial virus(Bartfeld & Clevers, 2015; Lawrence et al., 2024; van Dijk et al., 2024). These models have been useful in understanding immune responses, including those triggered by pathogens or during inflammatory conditions (Ali et al., 2024). Beyond structural and molecular hallmarks, we were able to leverage the aged mBEDO platform to provide a system for interrogating age-related mechanistic underpinnings of both immune-epithelial interactions and infectious diseases. Our platform demonstrates epithelial colonization and increasing bacterial burden after infecting aged mBEDOs with GFP-tagged UPEC, recapitulating previous reports on age-associated increase in recurrent UTIs *in vivo* (Ligon et al., 2023). mBEDOs amenability to co-culture with bone marrow–derived macrophages further amplify future applications of our platform to studying immune–epithelial interactions, which are often difficult to dissect *in vivo* due to the complexity of tissue microenvironments.

A key strength of the mBEDO platform is its adaptability for modeling genetic perturbations using tissues from genetic knockouts. We successfully generated mBEDOs from whole bladders of female *Irg1^−/−^* mice. Irg1 encodes a mitochondrial enzyme in the TCA cycle that catalyzes the conversion of cis-aconitate to itaconate, a metabolite with established immunomodulatory functions. While Irg1 has been extensively studied in myeloid cells, its role in epithelial aging remained unexplored. Loss of IRG1 resulted in metabolic profiles that closely resemble aged mBEDOs, particularly in purine and galactose metabolism. Notably, 35 out of 49 significantly altered metabolites in our three-way comparison followed a similar trend in aged and *Irg1^−/−^* mBEDOs, including increases in adenosine, guanine, and guanosine. These findings are consistent with prior reports of enhanced purine metabolism in Irg1-deficient macrophages and raise the possibility that loss of Irg1 activity may drive age-like metabolic shifts in bladder epithelial cells. Our data aligns with other studies that have shown itaconate is a key metabolite that offers a gero-protective role in inflammation and bone loss (Y. Wang et al., 2022). These data suggest that Irg1 plays a broader role in maintaining epithelial metabolic homeostasis and highlight the utility of mBEDOs for dissecting gene– metabolism interactions in the aging urothelium. Interestingly, metabolites linked to amino acid metabolism, such as D/L-serine and creatine, did not differ between *Irg1^−/−^* and young mBEDOs, suggesting that TCA cycle dysregulation in this context may selectively impact certain metabolic branches. In contrast, D-mannose, a galactose pathway metabolite and potential senotherapeutic, was reduced in both aged and *Irg1^−/−^* mBEDOs, while glucose-1-phosphate was increased. Nicotinamide, another age-dysregulated metabolite, was also significantly reduced in both groups. Together, these results support a model in which Irg1 contributes to metabolic homeostasis in the bladder epithelium and further highlights the mBEDO system as a powerful platform to study the intersection of metabolism, aging, and gene function in a tissue-specific context.

Organoids are emerging as invaluable tools for modeling metabolic signatures and exploring high-throughput metabolite supplementation(Murphy & Sweedler, 2022; Torrens-Mas et al., 2021). In recent years, metabolomics studies have been performed on *in vitro* organoid cultures in the context of pancreatic, kidney, as well as bladder cancer and infectious models(Ali et al., 2024; Keilberg et al., 2021; Walz et al., 2023), however, the targeted use of organoids specifically to address metabolic underpinnings of aging independent of disease has not been explored. We found that metabolites that were significantly increased in the aged samples could be mapped to the purine pathway with high confidence. Interestingly, purine metabolism has been reported to gradually increase with age in several models such as aged cardiac mouse tissue (Willems et al., 2003) and a dysregulated purine metabolism has been found to contribute to the progression of LUTS in the elderly (Birder & Jackson, 2021). In our study we found inosine, hypoxanthine, guanine, and adenosine were increased in aged mBEDOs. It has been previously reported that human fibroblast cultures from aged donors produced more inosine and adenosine *in vitro* that those from young donors (Ethier et al., 1989). The damage caused by oxidative stress to DNA may lead to the accumulation of 8-oxo-guanine lesions at telomeres, which would also explain the increase in guanine noted in aged mBEDOs with high oxidative stress and DNA damage (Barnes et al., 2022). A recent paper by Birder et al., has shown that exposure to hypoxanthine, which is known to increase oxidative stress, may exacerbate bladder aging as noted by voiding dysfunction and remodeling of the lower urinary tract (Birder et al., 2024). We also found phosphoethanolamine and choline levels were increased in aged mBEDOs. Accumulation of phosphoethanolamine has been implicated in senescence induction during aging, while elevated choline has been associated with increased incidence of overactive bladder symptoms, suggesting a link between lipid metabolism and functional decline in the aging bladder (Tighanimine et al., 2024). Previous studies have also reported non-neuronal production of choline in the bladder linked to higher incidences of OAB symptoms (Hanna-Mitchell et al., 2007). Identification of these specific pathways in our mBEDO platform lays the foundation to systematically test whether targeted interventions could ameliorate adverse metabolic changes with age.

Excitingly, we demonstrate supplementation with two age-dysregulated metabolites— Nicotinamide and D-mannose—restored redox balance and reduced DNA damage in aged mBEDOs. Both metabolites were significantly reduced in aged mBEDOs based on our metabolomics profiling. D-mannose, in particular, was previously shown by our group to exert gero-protective effects *in vivo* in the aged bladder epithelium (Joshi et al., 2024). Nicotinamide has also been widely studied for its role in lifespan extension and cellular resilience, largely due to its role as a precursor to nicotinamide adenine dinucleotide, a key cofactor in redox reactions and mitochondrial function (Anderson et al., 2003; Song et al., 2023). We observed that treatment with either Nicotinamide or D-mannose led to a marked reduction in oxidative stress and restoration of mitochondrial homeostasis, as evidenced by decreased ROS levels and lower mitochondrial mass. Reduction in intact mitochondrial abundance in aged mBEDOs may indicate improved mitochondrial function and adaptive response to energy demands (Srivastava, 2017). Previous studies show that reduction in mitochondrial abundance offers certain advantages such as improving metabolic flexibility and extending lifespan (Srivastava, 2017). Both interventions also significantly reduced γH2AX level, indicating a reduction in DNA damage in treated mBEDOs. These findings are consistent with prior work showing that Nicotinamide and its related precursors, such as Nicotinamide mononucleotide (NMN), can reduce oxidative stress and enhance DNA repair capacity in aging and disease models (John et al., 2012; Imai & Guarente, 2014). Several studies have shown NMN and Nicotinamide riboside which are both precursors of the same pathway help enhance DNA repair (Qiu et al., 2024). Similarly, D-mannose has recently been shown to reduce oxidative stress in other inflammatory conditions, including a murine model of ulcerative colitis (Lu et al., 2024), and reduction in osteoarthritis progression (Zhou et al., 2021) and its reduction in aged mBEDOs may contribute to a pro-senescent metabolic environment.

In sum, we have established murine bladder epithelium-derived organoids (mBEDOs) as a robust and scalable model that recapitulates key features of bladder aging, enables the study of epithelial–immune and host–pathogen interactions, and serves as a versatile platform for dissecting metabolic alterations and testing therapeutic interventions to restore epithelial homeostasis. Although our model does not yet incorporate the full complexity of bladder architecture, including vasculature and innervation, it offers a tractable and reproducible platform for studying epithelial aging. Future directions include extending this work to human bladder organoids derived from postmenopausal donors to reflect clinical relevance.

## Author Contributions

A.P., A.S., and I.U.M. conceived the experimental plan; A.P. performed the majority of experiments and was assisted by A.S., M.D., D.L., and A.S. Untargeted metabolomics and preliminary data analysis was performed by V.P. and N.P. S.J.B. analyzed the metabolomics data. A.P., A.S., and I.U.M. wrote the manuscript, and all authors approved the final draft.

## Acknowledgments

This work was supported in part by NIH grants, R56 AG084691-01A1 to I.U.M, and CPRIT grant through the Baylor College of Medicine Comprehensive Cancer Training Program under Award No. RP210027 to M.D. This research was also partially supported by NIH/NCI R01CA282282 and NIH U01CA111302 awarded to N.P. The metabolomics core was supported by the CPRIT Core Facility Support Award RP210227 “Proteomic and Metabolomic Core Facility,” and intramural funds from the Dan L. Duncan Cancer Center.

## Data Availability Statement

Data will be made available upon request.

## Additional Figures, Tables, and Legends

**Additional Table 1.**
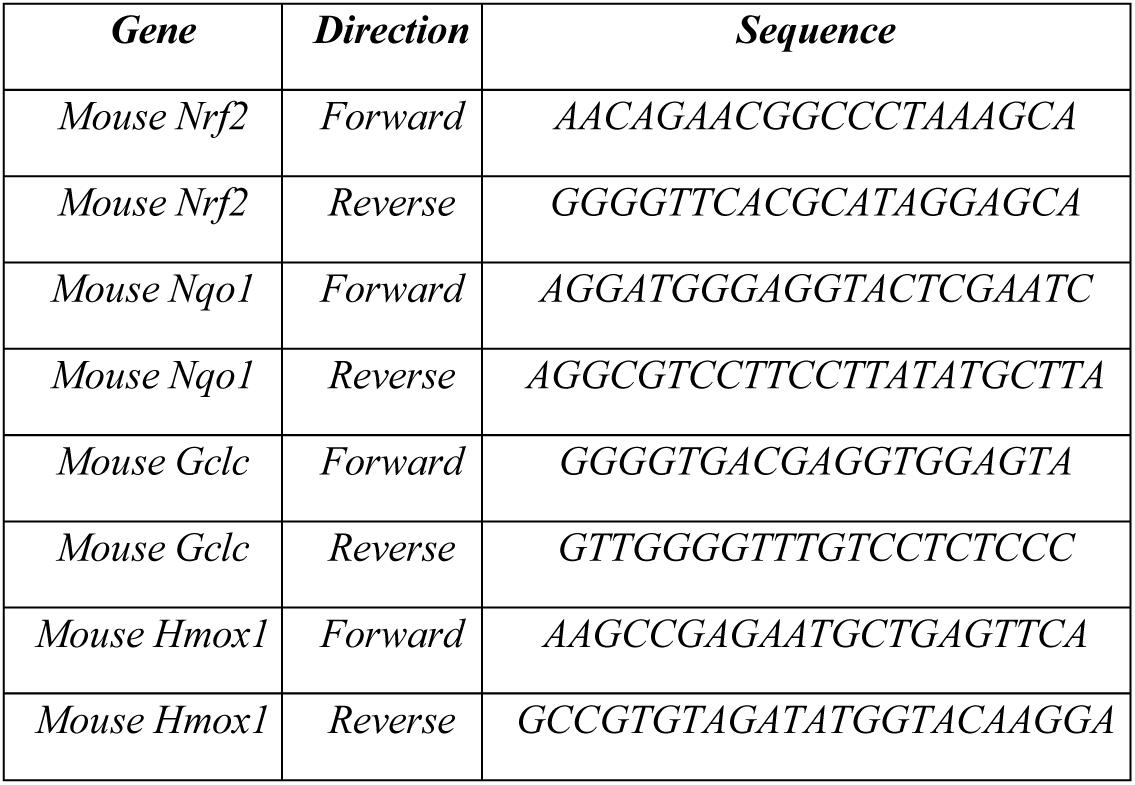
Primer information.

**Additional Figure 1.**
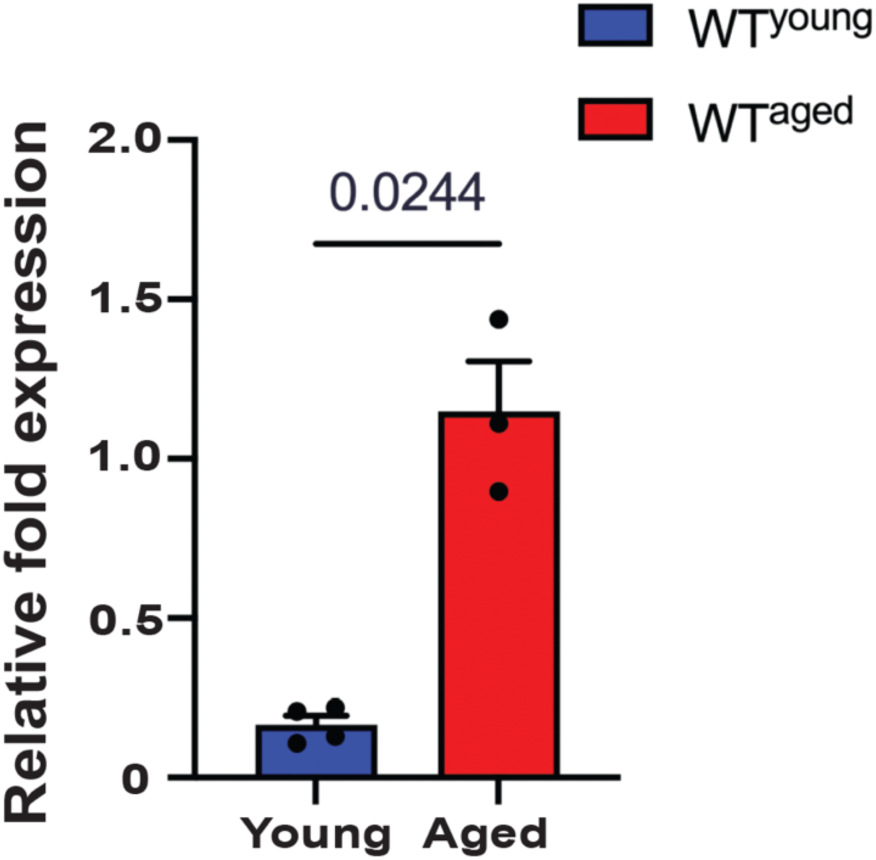
*Irg1* expression in young vs. aged mouse tissue. Relative fold change expression of *Irg1* in tissues of young and aged mice assessed via RNA Sequenci

